# Emerging SARS-CoV-2 variants reduce neutralization sensitivity to convalescent sera and monoclonal antibodies

**DOI:** 10.1101/2021.01.22.427749

**Authors:** Jie Hu, Pai Peng, Kai Wang, Bei-zhong Liu, Liang Fang, Fei-yang Luo, Ai-shun Jin, Ni Tang, Ai-long Huang

## Abstract

SARS-CoV-2 Spike-specific antibodies contribute the majority of the neutralizing activity in most convalescent human sera. Two SARS-CoV-2 variants, N501Y.V1 (also known as B.1.1.7 lineage or VOC-202012/01) and N501Y.V2 (B.1.351 lineage), reported from the United Kingdom and South Africa, contain several mutations in the receptor binding domain of Spike and are of particular concern. To address the infectivity and neutralization escape phenotypes potentially caused by these mutations, we used SARS-CoV-2 pseudovirus system to compare the viral infectivity, as well as the neutralization activities of convalescent sera and monoclonal antibodies (mAbs) against SARS-CoV-2 variants. Our results showed that N501Y Variant 1 and Variant 2 increase viral infectivity compared to the reference strain (wild-type, WT) *in vitro*. At 8 months after symptom onset, 17 serum samples of 20 participants (85%) retaining titers of ID_50_ >40 against WT pseudovirus, whereas the NAb titers of 8 samples (40%) and 18 samples (90%) decreased below the threshold against N501Y.V1 and N501Y.V2, respectively. In addition, both N501Y Variant 1 and Variant 2 reduced neutralization sensitivity to most (6/8) mAbs tested, while N501Y.V2 even abrogated neutralizing activity of two mAbs. Taken together the results suggest that N501Y.V1 and N501Y.V2 reduce neutralization sensitivity to some convalescent sera and mAbs.

## INTRODUCTION

Coronaviruses are enveloped, positive-stranded RNA viruses that contain the largest known RNA genomes to date. As severe acute respiratory syndrome coronavirus 2 (SARS-CoV-2) continues to circulate in the human population, multiple mutations accumulate over time, which may affect its transmission, virulence and antigenicity. Neutralizing antibodies (NAbs) elicited by natural infection or vaccination are likely to be a key immune correlate for protection against SARS-CoV-2 infection. Decline of antibodies response to SARS-CoV-2 in convalescent individuals and reinfections by different viral-variants have been reported ^1–3^. It is therefore important to gain insights into infectivity and antigenicity of SARS-CoV-2 variants.

Spike-specific antibodies contribute the majority of the neutralizing activity in most convalescent human sera. Two SARS-CoV-2 variants, N501Y.V1 (also known as B.1.1.7 lineage or VOC-202012/01) and N501Y.V2 (B.1.351 lineage), reported from the United Kingdom (UK) and South Africa, contain several mutations in the receptor binding domain (RBD) of Spike and are of particular concern. To address the infectivity and neutralization escape phenotypes potentially caused by these mutations, we used SARS-CoV-2 pseudovirus system to compare the viral infectivity, as well as the neutralization activities of convalescent sera and monoclonal antibodies (mAbs) against SARS-CoV-2 variants.

## METHODS

The blood samples (n = 40) of 20 patients with COVID-19 obtained in February and October 2020 in Chongqing were previously reported.^2^ Eight RBD-specific mAbs with neutralizing capability against SARS-CoV-2 were obtained from the blood samples of COVID-19 convalescent patients.^4^ DNA sequences encoding reference strain (wild-type, WT) and mutant Spike proteins of SARS-CoV-2 were codon-optimized and synthesized by Sino Biological Inc (Beijing, China) and GenScript Inc (Nanjing, China). Using luciferase-expressing lentiviral pseudotype system, we expressed WT, N501Y.V1 (Variant 1) and N501Y.V2 (Variant 2) mutant Spike proteins in enveloped virions, respectively. The neutralizing antibodies (NAbs) were measured by pseudovirus-based assays in 293T-ACE2 cells. The inhibitory dose (ID_50_) was calculated as the titers of NAbs.

### Ethical Approval

The study was approved by the Ethics Commission of Chongqing Medical University (reference number 2020003). Written informed consent was waived by the Ethics Commission of the designated hospital for emerging infectious diseases.

## RESULTS

First, the infectivity of pseudotyped viral particles were measured by luciferase assay as previously described.^5^ As shown in Fig.1a, the entry efficiencies of Spike pseudotyped viruses bearing N501Y Variant 1 or Variant 2 mutant were about 3 to 4.4 times higher than that of the WT pseudovirus when viral input was normalized, suggesting that these spike variants promote the infectivity of SARS-CoV-2. Then, we assessed the neutralizing efficacy of 40 convalescent serum samples from 20 individuals at two time points with pseudovirus neutralization assay. At follow-up time point 1, corresponding to a median of 25 days (range 5–33 days) post-symptom onset, most sera were significantly less effective in neutralizing the N501Y Variant 1 and Variant 2 compare to WT pseudovirus (Fig. 1b). The mean nAb titers were 825 for WT, 343 for Variant 1, and 148 for Variant 2. The neutralizing activity of 2 samples against N501Y.V1 was reduced by >10-fold. Notably, the NAb titers of 6 samples (30%) decreased below the threshold against Variant 2 (Fig. 1b). At follow-up time point 2 (about 8 months post-symptom onset), 17 samples of 20 participants (85%) retaining titers of ID_50_ >40 against WT pseudovirus, whereas the NAb titers of 8 samples (40%) and 18 samples (90%) decreased below the threshold against N501Y Variant 1 and Variant 2, respectively (Fig. 1c). These data indicate that N501Y Variant 1 and Variant 2 escape from neutralizing antibodies in some COVID-19 convalescent sera.

**Fig.1.**
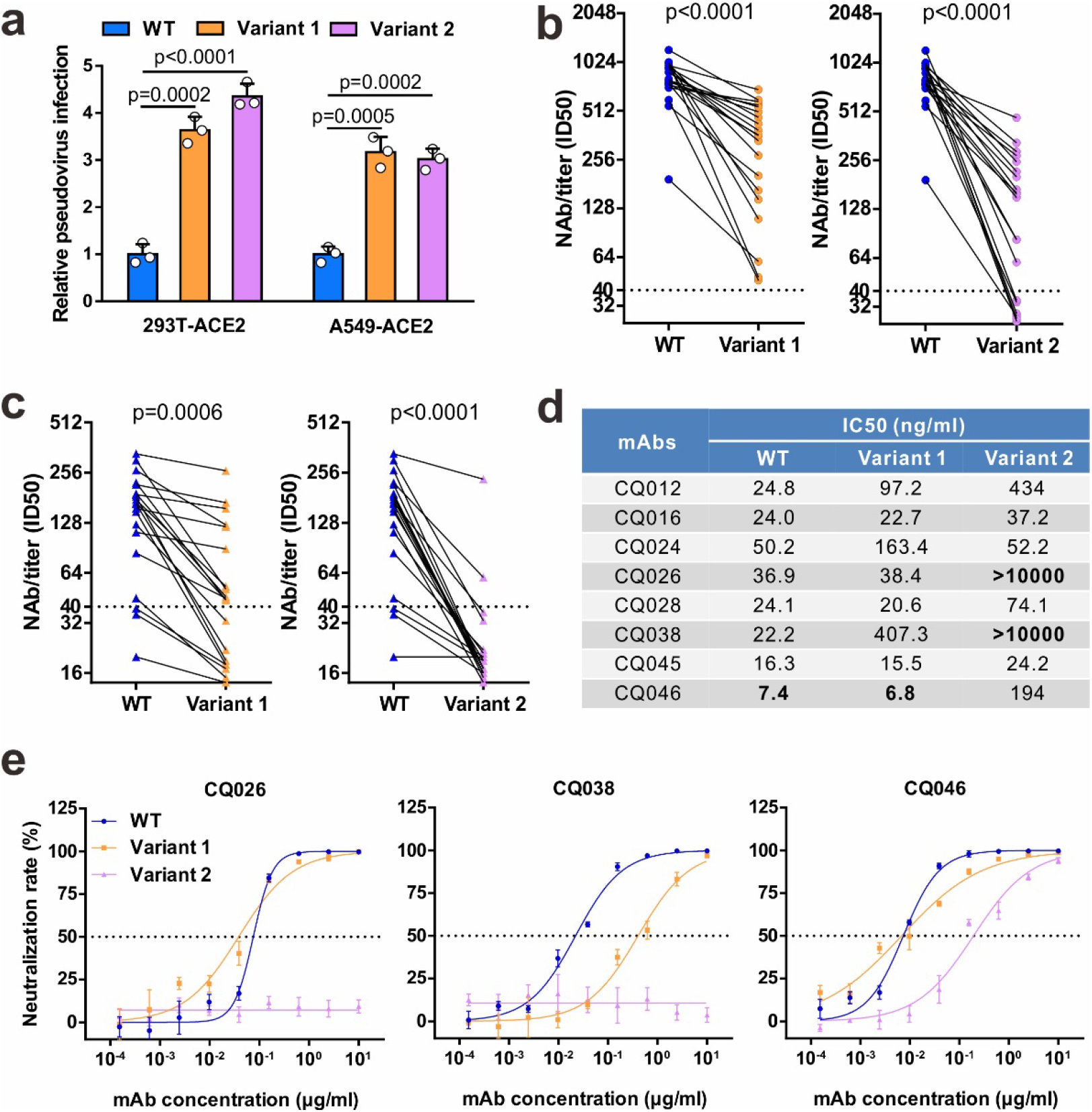
Neutralizing activities of convalescent sera and monoclonal antibodies against SARS-CoV-2 variants. **a** Infectivity of WT and variant pseudovirus conducted in 293T-ACE2 and A549-ACE2 cells. Cells were inoculated with equivalent doses of each pseudotyped virus. WT, wild-type Spike (GenBank: QHD43416) pesudotyped virus; Variant 1, N501Y.V1 mutant Spike pesudotyped virus (containing H60/V70 deletion, Y144 deletion, N501Y, A570D, D614G, P681H, T716I, S982A, D1118H); Variant 2, N501Y.V2 mutant Spike pesudotyped virus (containing K417N, E484K, N501Y, D614G). **b-c** Neutralization of WT and variant pesueoviruses by convalescent sera. Pseudovirus-based neutralizing assay were performed to detect neutralizing antibody (NAb) titers against SARS-CoV-2. The thresholds of detection were 1:40 of ID50. Twenty sera (indicated by circles) were drawn 5 to 33 days post-symptom onset (**b**); 20 sera (indicated by triangles) were drawn ~ 8 months post-symptom onset (**c**). **d-e** The half-maximal inhibitory concentrations (IC50) for tested monoclonal antibodies (mAbs) against pseudoviruses (**d**) and representative neutralization curves (**e**). Statistical significance was determined by One-way ANOVA.

In addition, we assessed the impact of these variants on neutralizing activity of human monoclonal antibodies (mAbs) isolated from COVID-19 convalescent patients. All eight antibodies potently neutralized the WT pseudovirus, while two mAbs (CQ016 and CQ045) are only minimally affected by the variants. However, the neutralization activities of six mAbs were reduced or abolished by either N501Y Variant 1 or Variant 2 (Fig. 1d). Among them, three mAbs were less effective against N501Y.V1 and five against N501Y.V2 by 3-folds or more (Fig. 1d). Notably, two mAbs (CQ026 and CQ038) showed no neutralizing activity to N501Y.V2. Moreover, the Variant 2 reduced the neutralization sensitivity with the most potent mAb CQ046 by 26 folds, compared with that of WT pseudovirus. The IC_50_ of mAb CQ046 increased from 7.4 ng/ml (WT) to 194 ng/ml (Variant 2) (Fig. 1e). Together, both N501Y Variant 1 and Variant 2 reduced neutralization sensitivity to most mAbs tested, while N501Y.V2 even abrogated neutralizing activity of two mAbs.

## DISCUSSION

Our findings indicated that N501Y Variant 1 and Variant 2 increase viral infectivity compared to the reference strain *in vitro*. Notably, both N501Y Variant 1 and Variant 2 contain the D614G and N501Y mutations in Spike protein. The findings that Variant 1 and Variant 2 enhanced the infectivity of SARS-CoV-2 *in vitro* are highly consistent with previous studies, which demonstrated that D614G and N501Y mutations enhanced the fitness and transmissibility of the virus as evidenced by structure analysis and the increased number of clinical cases.^6,7^ Another key question is whether some mutations may enable immune evasion. It is reported that neutralization escape mutants can be selected by passaging virus in the presence of NAbs.^8^ Here, we observed that two naturally occurring SARS-CoV-2 variants, N501Y Variant 1 and Variant 2, were more resistant to neutralization by some mAb and convalescent sera from patients that were infected in mid- to late- January 2020 when a ‘first wave’ virus was mainly circulating in China. Consistently, Spike variants with the H60/V70 deletion or E484K mutation have significantly reduced susceptibility to neutralization by the polyclonal serum antibodies of some individuals.^9,10^ Whether these patients were at high risk of reinfection with ‘second wave’ variants should be explored in further studies. It is also urgent to assess the effectiveness of currently authorized vaccines against these variants.

Collectively, this study will be helpful for understanding SARS-CoV-2 infectivity and for the design of vaccines against COVID-19. Given the evolving nature of the SARS-CoV-2 RNA genome, antibody therapeutics and vaccine development require further considerations to accommodate mutations in Spike that may affect the antigenicity of the virus. Limitations of this study include its small sample size and the use of non-replicating pseudovirus system. Therefore, further studies with authentic SARS-CoV-2 viruses are required.

## Acknowledgements

We would like to thank Prof. Cheguo Cai (Wuhan University, Wuhan, China) for providing the pNL4-3.Luc.R-E- lentiviral plasmid. Our work has received funding support from the Emergency Project from the Science & Technology Commission of Chongqing (cstc2020jscx-dxwtB0050, cstc2020jscx-fyzx0053), the Emergency Project for Novel Coronavirus Pneumonia from the Chongqing Medical University (CQMUNCP0302), the Key Laboratory of Infectious Diseases (CQMU, 202005), the Leading Talent Program of CQ CSTC (CSTCCXLJRC201719), and a Major National Science & Technology Program grant (2017ZX10202203) from the Science & Technology Commission of China.

## Author Contributions

A.H., N.T., and A.J. developed the conceptual ideas and designed the study. J.H. and P.P. performed the experiments. B.L. and L.F. provided the samples. F.L. was responsible for mAb purification. K.W. performed statistical analysis. All authors provided scientific expertise and the interpretation of data for the work. K.W. drafted the manuscript. All authors have approved the final version of the manuscript.

